# Belowground plant responses to root herbivory depend on the composition and structure of their root-colonising arbuscular mycorrhizal fungi

**DOI:** 10.1101/2022.02.28.480478

**Authors:** Anna Ng, Bree A.L. Wilson, Adam Frew

## Abstract

1. There is growing interest in managing arbuscular mycorrhizal (AM) fungi in agriculture to support plant production. These fungi can support crop growth and nutrient uptake but also affect plant-herbivore interactions. Our knowledge of how native AM fungal diversity and community composition influence these interactions is limited, while our understanding of this in relation to root-herbivory is lacking altogether.
2. To begin to address these knowledge gaps, plants were grown with no AM fungi or were inoculated with native fungal communities sourced from either a crop field (field community), a sclerophyll forest (forest community), or a crop field in fallow (fallow community). We then explored how the composition and structure (species richness and relative abundance) of root-colonising AM fungal communities was associated with the growth and belowground nutrient responses of a major crop (*Sorghum bicolor*) to attack from a root-feeding insect (*Dermolepida albohirtum*).
3. DNA metabarcoding revealed plants associated with three distinct root-colonising AM fungal communities. Fungal taxon richness in roots was highest in the field community and lowest in the fallow community. Both the field and fallow communities were dominated by the putatively ruderal genera *Glomus* and *Claroideoglomus*, while the forest-derived community contained greater proportions of *Paraglomus* and *Ambispora*.
4. In response to root herbivory, plants without AM fungi and plants colonised by the forest community exhibited root biomass losses of 61% and 44%, respectively. Similarly, these two groups also had reductions of 59% and 65% in their root phosphorus content, respectively, when subjected to the root herbivore. In contrast, plants associating with communities harbouring greater proportions of *Glomus* and *Claroideoglomus* (the field and fallow communities) did not exhibit reductions in root biomass or nutrient content.
5. Our results show that plant responses to root-herbivory vary with root-colonising AM fungal community composition and structure. In a community context, our findings suggest that stronger associations with the genera *Claroideoglomus* and *Glomus* may potentially support crop tolerance-associated responses belowground. There is an urgent need for more exploration of how natural assemblages of AM fungi differentially mediate plant-herbivore interactions if we are to effectively manage soil fungi in sustainable agricultural systems.

## INTRODUCTION

The importance of belowground interactions of plants with other soil biota is being increasingly recognised in both natural and managed ecosystems (Bardgett & van der Putten, 2014), including in agriculture (Bender *et al.*, 2016). As part of this, a key goal is the effective management of soil biota to enhance soil ecosystem function and the resultant ecosystem services to support agricultural production. Arbuscular mycorrhizal (AM) fungi are a group of soil and plant dwelling fungi (subphylum Glomeromycotina; Spatafora *et al.*, 2016) that are a central component of improving sustainable management of soil biota in crop systems (Rillig *et al.*, 2019). These fungi are globally widespread and are reliant on symbiotic interactions with plants for their carbon, while providing their plant hosts with access to resources in the soil including nutrients (primarily phosphorus) and water (Smith & Read, 2008). In the context of plant production, AM fungi are typically considered for their potential to enhance plant growth, nutrient uptake, and yield. Yet they play important roles in various ecosystem processes such as nutrient cycling, decomposition, soil aggregation, and plant-plant competition, hence their heralded importance to soil sustainability (Powell & Rillig, 2018; Tedersoo *et al.*, 2020).

AM fungi (AMF) also indirectly affect plant fitness by mediating plant interactions with insect herbivores (Hartley & Gange, 2009; Vannette & Rasmann, 2012; Borowicz, 2013). To deal with insect herbivory, plants can rely on resistance-associated defence, which negatively impacts herbivore performance (e.g., growth, fecundity, survival) or preference for the host plant. Additionally, AM fungi can also affect plant tolerance-based responses to herbivory, which is the ability for the plant to regrow (compensatory growth) and reproduce after herbivore attack. This is not unexpected considering AM fungi can improve plant nutrient uptake, which then may better equip them to tolerate tissue loss. Yet the outcomes for plant tolerance are quite variable and are not clearly linked to particular plant groups, or herbivore feeding guilds (Borowicz, 2013).

Compared to our knowledge on AM fungal-plant-herbivore interactions aboveground, our understanding of these interactions involving belowground herbivores is limited (Johnson & Rasmann, 2015). Root herbivores are often negatively impacted by the AM symbiosis, an outcome that may be driven by changes in silicon-based defences (Frew *et al.*, 2017), or other defence-associated chemistry (Barber *et al.*, 2013; Hill *et al.*, 2018). Research has also shown that increases in aboveground plant growth, as a form of tolerance to rootherbivory, can be enhanced by the AM symbiosis (Frew *et al.*, 2020).

The consequences of these interactions for plants not only depend on plant and herbivore identities, but also on the identities of the root-colonising AM fungi (Bennett & Bever, 2007; Tao *et al.*, 2016). The functional diversity of AM fungi is widely acknowledged (Van Der Heijden & Scheublin, 2007; Powell & Rillig, 2018), but is often overlooked in the context of herbivory (Frew *et al.*, 2022). Many studies have shown that different AM fungal taxa have differing impacts on plant responses to herbivory, including tolerance (Bennett & Bever, 2007; Borowicz, 2013) and resistance (Wooley & Paine, 2007; Vannette *et al.*, 2013; He *et al.*, 2017). Furthermore, fungal species richness can also shape these relationships, for example, Bennet *et al.* (2009) found the taxon-specific effects of two AM fungi were lost if they were inoculated as a mixed community. Contrastingly, recent work found plant defence responses to aboveground herbivory increased with fungal species richness (Frew & Wilson, 2021). However, most research to date has focussed on a small handful of cosmopolitan fungal taxa (e.g. *Rhizophagus irregularis, Funneliformis mosseae;* Malik, 2018). There are around 288 described species of AM fungi (c. 1700 putative species; Öpik & Davison, 2016), and most plants, including crop species, associate with multiple AM fungal species concurrently in the field (Öpik *et al.*, 2013; Bainard *et al.*, 2014). Continuing to focus on a few fungal taxa, or manipulated assemblies of commonly-used taxa, has perpetuated a narrow understanding of AM fungal-plant-herbivore ecology (Frew *et al.*, 2022). This represents an important barrier in our ability to incorporate the management of AM fungi to agriculture. For example, the application of ‘foreign’ fungal taxa as inoculants has shown limited success in the field (see examples in Hart *et al.*, 2018) which is unsurprising as the underlying assumption is that the introduction of fungi to an established soil fungal community will result in positive plant outcomes (Rodriguez & Sanders, 2015).

A handful of studies have explored how naturally occurring fungal communities affect interactions between plants and insect herbivores. For example, Kula *et al.* (2005) examined plant responses to aboveground herbivory with and without colonisation by a native AM fungal community, where the authors found that the fungi enhanced plant growth in response to herbivore attack. Barber *et al.* (2013) investigated belowground plant trait responses using a commercial single species inoculum and two (unidentified) native fungal communities. Here the authors found community-specific effects on plant nutrients and on belowground defence chemistry. However, such studies neglect to identify the fungal communities in the roots. Despite continued interest in the potential of using AM fungi in agriculture (Thirkell *et al.*, 2017; Hart *et al.*, 2018), our understanding of how different native communities of AM fungi can affect plant responses to herbivory is limited (Frew *et al.*, 2022), and this knowledge in the context of belowground herbivory is effectively absent.

To begin to address these knowledge gaps, this study explored how inoculation with different native AM fungal communities, and the resultant differences in the composition and structure (species richness and relative abundances) of the root-colonising communities, affected plant responses to attack from a root-feeding insect. To do this we used sorghum (*Sorghum bicolor*), an important crop species which forms arbuscular mycorrhizas. Root-herbivory was applied using the canegrub (*Dermolepida albohirtum*), a native insect to Australia and a significant root-feeding pest of various crops (Allsopp, 2010; Sallam, 2011). Plants were grown with no AM fungi, or they associated with one of three different fungal communities that were sourced from three adjacent locations in southeast Queensland, Australia. The three locations were selected as they were under different management such that we expected distinct differences in community composition (Moora *et al.*, 2014). We hypothesised that: (i) AM fungal alpha diversity in plant roots would be positively associated with greater tolerance to root-herbivory (i.e. belowground plant growth and nutrient status would be less negatively affected by root-herbivory); (ii) plant responses to root-herbivory would vary with differences in community composition and abundance of root-colonising AM fungi.

## METHODS

We conducted a factorial pot experiment with two factors of ‘AMF’ and ‘Herbivore’ using 64 individual sorghum plants (*Sorghum bicolor* L. Moench cv. “MR Taurus”). Plants associated with one of three different soil-derived AM fungal communities, or their roots were absent from AM fungi. To achieve this, prior to seed germination, 10 L pots were filled with 1:1 sand (Ki-Carma® Landscape sand) and soil (Richgro® All Purpose Garden Soil) which had been pasteurised at 80 °C for 2 h. The different native AM fungal inocula treatments were sourced from three sampling locations within the same 5 km^2^ area of southeast Queensland, Australia (−27.4326°, 152.3495°). These three community inocula were: (i) ‘Field community’ soil which was sourced from an organically managed arable field with recently harvested sweetpotato (*Ipomoea batatas* ‘Bellevue’); (ii) ‘Forest community’ soil, which was sourced from an adjacent remanent sclerophyll forest with a mixed community of *Eucalyptus* spp., *Acacia* spp., *Allocasauarina* spp., and a mixed understory of native grasses, and; (iii) ‘Fallow community’ soil, which was sourced from a neighbouring arable field which had been in fallow for 12 months and previously harboured a crop of snow peas (*Pisum sativum*). At each of these sites, soil was taken from five randomly chosen 1m x 1m plots and then pooled for the experiment. Each of these soil inocula (200 ml) were homogenised, sieved, and then added to the top 20 cm of the pots and mixed through (16 pots each), while the no AMF pots (n=16) received 200 ml of equal parts from all three soils, sterilised at 121°C for 20 mins. As the addition of the whole soil inocula introduces other soil microbes along with AM fungi, all 64 pots received 250 ml of a microbial wash to standardise the non-AM fungal microbial community. This wash was composed of the filtrate from a washed soil mix made from equal parts of the three soils filtered through a 38 μm sieve (Koide & Li, 1989). At this stage a 25 g sample of soil from the top 20 cm was taken from four of each of the AMF treatments for soil nutrient analyses (Table S1).

Sorghum seeds were initially germinated in Petri dishes (90 mm ⍰) with water for 7 days after which a seedling was carefully planted into the top 5cm of each pot. Plants were watered as necessary and were randomly re-distributed throughout the glasshouse chamber every week to avoid any spatial effects. Air temperature was regulated with a 14 h day: 10 h night mean of 27 °C: 17 °C (± 3 °C). After 14 weeks, half of the pots under each of the four treatments were subjected to the root herbivore treatment where a third-instar *Dermolepida albohirtum* larva was weighed then placed in the top 5 cm layer of soil. These insects are scarab beetles native to Australia, the larvae are generalist herbivores that feed of the roots of grasses (Sallam, 2011). Larvae were supplied by Dr Pauline Lenancker (Sugar Research Australia Pty Ltd) and were reared exclusively on carrot prior to the experiment. After two weeks, plants and root herbivores were removed from their pots. Root herbivores were weighed and the relative growth rate (RGR) was calculated as per Slansky (1985). The above- and belowground plant material were separated, roots were washed and subsamples of roots were taken from each plant for subsequent assessment of fungal root colonisation. The remaining plant material was oven dried at 40 °C for 72 h, then weighed. Belowground tissue was ground to a fine powder and homogenised prior to chemical analysis.

### AM fungal colonisation scoring

To confirm colonisation of roots inoculated with the three AM fungal soils and the absence of colonisation in the plants under the no AMF treatment, the 1-2g root subsample was cleared with 10% potassium hydroxide heated to 80 °C for 20 mins and then stained with 5% ink vinegar (Vierheilig *et al.*, 1998). The cleared and stained roots were mounted on glass slides with glycerine under a cover slip and scored for the presence of AM fungi using the intersect method (McGonigle *et al.*, 1990). To conservatively quantify colonisation, only hyphae with a clearly visible connection to AM fungal structure (i.e., arbuscule, vesicle, spore) were counted.

### Root chemistry

To measure root carbon and nitrogen, dried and ground root material was analysed using the high temperature combustion method (LECO analyser) where samples are loaded into a combustion tube and flushed with oxygen. Gases generated from this process are then measured using an infra-red detector for carbon, and a thermal conductivity cell for nitrogen. Root phosphorus, potassium, sulphur, copper, zinc, manganese, calcium, magnesium, sodium, and boron were measured using inductively coupled plasma (ICP) spectroscopy after digestion of the dried root material with hydrogen peroxide and nitric acid (Rayment & Lyons, 2011). Plant nutrient analyses were carried out by CSBP Soil and Plant Analysis Laboratory (Bibra Lake, Western Australia).

### DNA extraction, sequencing, and bioinformatics

DNA was extracted from 70 mg of dried root samples using a DNeasy Powersoil Pro Kit (Qiagen, GmBH, Hilden, Germany) according to the manufacturer’s instructions, with the modification that dried root material cut into small 0.5 mm fragments is added to extraction tubes instead of soil. Sequencing was processed through the Western Sydney University’s Next-Generation Sequencing Facility’s (Richmond, NSW, Australia) liquid handling pipeline using in-house optimised protocols. The DNA was purified using the Agencourt AMPure XP Beads (Beckman Coulter), followed by a quality assessment using the Qunat-iT ™ PicoGreen fluorescence-based analysis (ThermoFisher Scientific, North Ryde, NSW, Australia). The purified DNA then underwent amplification using polymerase chain reaction (PCR) targeting the small-subunit (SSU) ribosomal RNA gene using AM fungal-specific primers: WANDA (Dumbrell *et al.*, 2011) and AML2 (Lee *et al.*, 2008). Briefly, PCR reactions consisted of 5μl Q5® High-Fidelity DNA Polymerase 2 x master mix (New England Biolabs, Notting Hill, Victoria, Australia), 0.2 μl of 10 μM forward and reverse primer with 2 μl of DNA, total volume of the reaction was 10 μl. PCR reactions were as described in Caporaso et al. (2018). The amplified PCR product then underwent a short second PCR to attach the Illumina Nextera XT v2 index set (Illumina Australia, Melbourne, Australia), as per manufacturer’s instructions (https://support.illumina.com). Each reaction consisted of 3.8 μl of Q5® High-Fidelity DNA Polymerase 2x master mix (NEB), 1.5 μl of each index (Illumina) and 2.3 μl of PCR product, total reaction volume was 7.6μl. The amplicons were then diluted and again assessed using the Qunat-iT™ PicoGreen (ThermoFisher Scientific) assay and normalised library pools were constructed using the Eppendorf epMotion. The libraries were cleaned-up and prepared for sequencing following the Illumina MiSeq protocol. Sequencing was performed on the Illumina MiSeq platform using the Illumina MiSeq reagent kit v3 2x 300bps paired-end chemistry as per manufacturer’s instructions.

Bioinformatic data analysis and processing was conducted using the graphical downstream analysis tool (gDAT) for analysing rDNA sequences (Vasar *et al.*, 2021). Raw reads (20⍰×⍰2,412,166 reads in total) were demultiplexed and cleaned using a series of bioinformatic steps (see Vasar *et al.*, 2017, 2021). In short, reads were demultiplexed by checking double barcodes, allowing one mismatch for both reads. Reads were retained if they carried the correct primer sequences (WANDA and AML2; allowing one mismatch for each) and had an average quality of at least 30, and orphan reads were removed (leaving 20⍰×⍰1,672,238 cleaned reads). Putative chimeric sequences (7,636; 0.5% of cleaned reads) were identified and removed using vsearch v2.15.0 (Rognes *et al.*, 2016) with the default parameters in reference database mode against the Maarj*AM* database (status July 2021; Öpik *et al.*, 2010). Cleaned and chimera-free sequences were assigned to virtual taxa (VT) using BLAST+ (v2.7.1, Camacho et al. 2009) referencing the Maarj*AM* database (Öpik *et al.*, 2010) with at least 97% identity and 95% alignment. Non-AM fungal sequences were identified with a BLAST+ search against GenBank, most of these non-AM fungal sequences were assigned to plants (87%), metazoa (6%), and fungi (3%).

### Statistics

All analyses were carried out using R statistical interface v4.0.5 (R Core Team, 2017) and RStudio v1.4.1717.

To examine how different native communities of AM fungi affected plant growth and nutrient responses to root-herbivory, we first analysed and compared the root-colonising fungal communities assembled in response to the initial inoculations and also assessed if they were affected by root-herbivory. The ‘no AMF’ plant roots were not sequenced (root staining confirmed absence of colonisation), and thus not included in the fungal community analyses.

To counteract bias from differences in sequencing depth, samples were rarefied to an even depth by means of the *rarefy_even_depth* function from the R package ‘Phyloseq’ (McMurdie & Holmes, 2013). Alpha diversity of AM fungi under each treatment combination was assessed as observed virtual taxon (VT) richness. The effects of the AMF and herbivore treatments on VT richness were assessed by fitting standard linear models using the *lm* function comparing the two factors ‘AMF’, ‘Herbivory’ and their interaction, then applied *Anova* function from the R package ‘car’ (Fox & Weisberg, 2011). Dissimilarity in community composition and structure of the root-colonising AM fungal communities were examined using principle coordinate analysis (PCoA, package ‘Phyloseq’; McMurdie & Holmes, 2013) based on Bray-Curtis dissimilarity. The effects of the AMF and herbivore treatments were analysed using permutational multivariate ANOVA (perMANOVA) with post-hoc analysis comparing AM fungal groupings in the PCoA using the *adonis* function from the R package ‘vegan’ (Oksanen *et al.*, 2015), and the *perwise.perm.manova* function from the R package ‘RVAideMemoire’ (Hervé & Hervé, 2020). To explore preferential treatment-taxa associations at the VT and genus level of taxonomic resolution, we used indicator species analysis from the R package ‘indicspecies’ (Cáceres & Legendre, 2009). To further explore differences in community composition between the three root-colonising communities, the relative abundances of AM fungal genera was analysed by fitting generalised linear models (family = quasibinomial) using the *glm* function and applied the *Anova* function from R package ‘car’.

To assess how the different AM fungal communities and root-herbivory affected plant growth and root nutrient content, we fitted standard linear models using the *lm* function and applied the *Anova* function from ‘car’ package in R. Data exploration was carried out following the protocol described in Zuur *et al.* (2010). Root content of phosphorus, zinc, manganese, nitrogen, sulphur, and calcium response variables were all log transformed to give residual diagnostic plots which fit a normal distribution. To investigate any differences in AM fungal root colonisation under the different treatments, we fitted negative binomial generalised linear models using the *glm.nb* function from the R package ‘MASS’ (Venables & Ripley, 2002). To determine if there were any differences in root-herbivore performance (relative growth rate) while feeding on plant roots colonised by the different AM fungal communities, or with no AM fungi, we fitted a standard linear model and applied the *Anova* function.

## RESULTS

### AM fungal diversity and community composition

High-throughput sequencing of AM fungal DNA present in the root samples yielded amplicons identified as known AM fungal species (virtual taxa or VT). After rarefaction to normalise sequence data, a total of 34 VT were recorded across all root samples **(Table S2).** The four most dominant taxa in the dataset made up 73.9% of all AM fungal sequences and were members of the genera *Claroideoglomus* (VT193; 35.8%), *Paraglomus* (VT444; 17.8%) and *Glomus* (VT67; 8.9%, and VT92; 9.8%; Fig. 1). Indicator species analysis revealed particular VT were strongly associated with the root-colonising AM fungal communities derived from the field, forest, and fallow soil inoculations, these root communities are hereafter referred to as the ‘field community’, ‘forest community’, and ‘fallow community’. AM fungal VT67 (*Glomus* sp.) was strongly associated with the field community, four VT (comprising *Ambispora* sp., *Glomus* sp., *Claroideoglomus* sp., and a *Scutellospora* sp.) were indicators of the forest community, while two VT (VT279 and VT57; *Claroideoglomus* spp.) were indicator VT of the fallow community **(Table 1**). Further analysis revealed preferential associations of particular AM fungal genera with particular communities. Specifically, *Ambispora, Paraglomus*, and *Scutellospora* were all indicator genera of the forest community, *Claroideoglomus* was an indicator of the fallow community, while *Glomus* was particularly associated with both the field and fallow communities **(Table 1**). There were significant differences in the relative abundances of different AM fungal genera between the three root-colonising fungal communities **(Fig. 2).** Specifically, *Glomus* was most abundant in the field community **(Fig. 2a),** while *Claroideoglomus* had the highest abundance within the fallow community **(Fig. 2b).** The community derived from the forest soil inoculation held the highest relative abundances of both *Paraglomus* **(Fig. 2c)** and *Ambispora* **(Fig. 2d).**

**Fig. 1.**
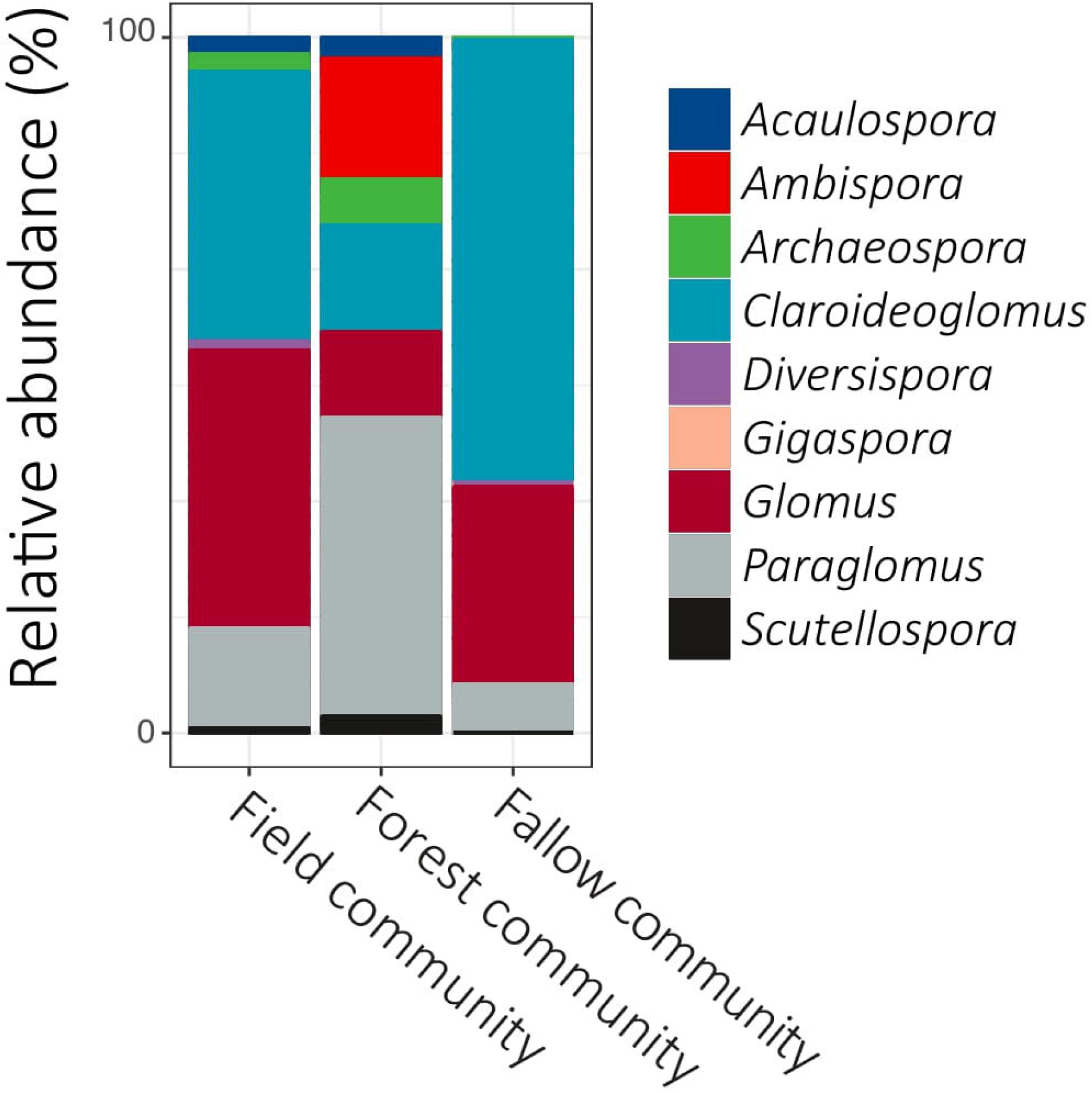
Relative abundance of root-colonising arbuscular mycorrhizal (AM) fungal genera from plants inoculated with one of three different AM fungal communities sourced from a cropped agricultural field (‘field community’), a sclerophyll forest (‘forest community’), or a cropped agricultural field in fallow (‘fallow community’).

**Fig. 2.**
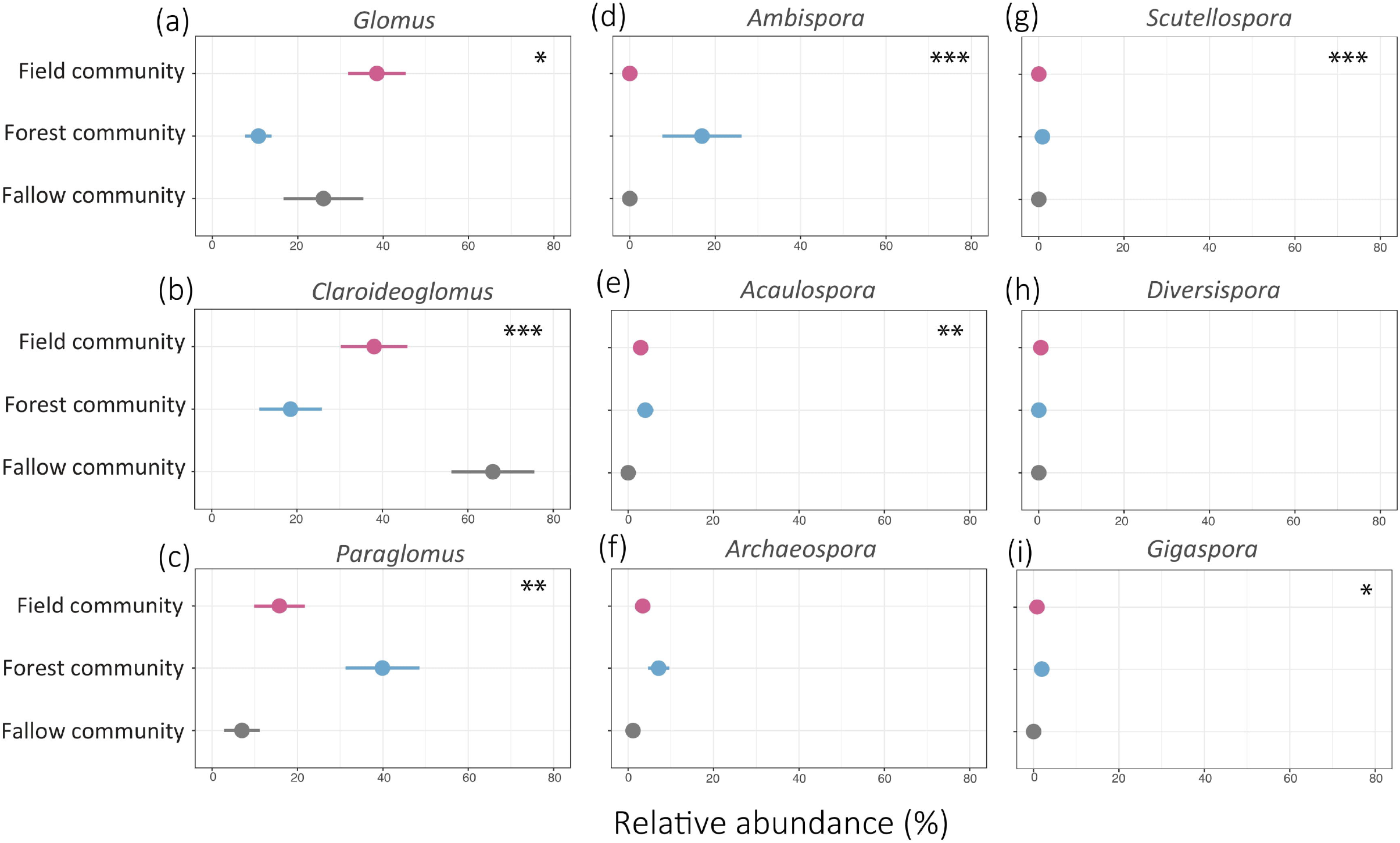
The relative abundances of arbuscular mycorrhizal (AM) fungal genera **(a)** *Glomus*, **(b)** *Claroideoglomus*, **(c)** *Paraglomus*, **(d)** *Ambispora*, **(e)** *Acaulospora*, **(f)** *Archaeospora*, **(g)** *Scutellospora*, **(h)** *Diversispora*, and **(i)** *Gigaspora* colonising roots of plants inoculated with three different AM fungal communities sourced from a cropped agricultural field, a sclerophyll forest, or a cropped agricultural field in fallow. Significant effects of the AM fungal community treatment are indicated * *P* <0.05, ** P<0.01, *** *P* <0.001, values are means ± SE

**Table 1.**
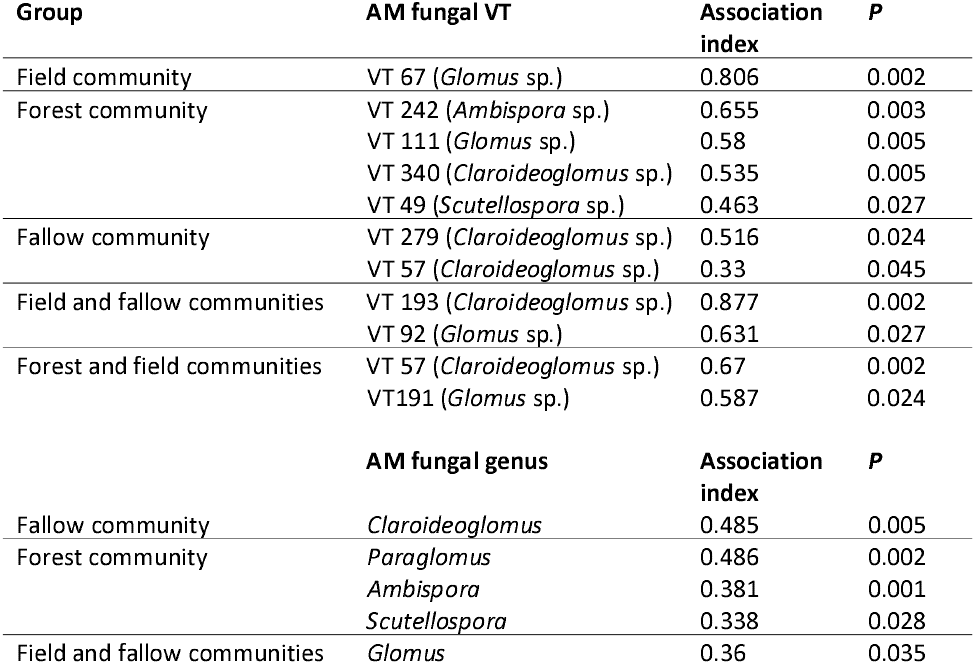
Indicator virtual taxa (VT) and genera of arbuscular mycorrhizal (AM) fungi for different root-colonising fungal communities in plant (*Sorghum bicolor*) roots. Values obtained from the *multipatt* function including the association index and associated *P*-value.

| Group | AM fungal VT | Association index | *P* |
| --- | --- | --- | --- |
| Field community | VT 67 (*Glomus* sp.) | 0.806 | 0.002 |
| Forest community | VT 242 (*Ambispora* sp.) | 0.655 | 0.003 |
|  | VT 111 (*Glomus* sp.) | 0.58 | 0.005 |
|  | VT 340 (*Claroideoglomus* sp.) | 0.535 | 0.005 |
|  | VT 49 (*Scutellospora* sp.) | 0.463 | 0.027 |
| Fallow community | VT 279 (*Claroideoglomus* sp.) | 0.516 | 0.024 |
|  | VT 57 (*Claroideoglomus* sp.) | 0.33 | 0.045 |
| Field and fallow communities | VT 193 (*Claroideoglomus* sp.) | 0.877 | 0.002 |
|  | VT 92 (*Glomus* sp.) | 0.631 | 0.027 |
| Forest and field communities | VT 57 (*Claroideoglomus* sp.) | 0.67 | 0.002 |
|  | VT191 (*Glomus* sp.) | 0.587 | 0.024 |
|  | **AM fungal genus** | **Association index** | ***P*** |
| Fallow community | *Claroideoglomus* | 0.485 | 0.005 |
| Forest community | *Paraglomus* | 0.486 | 0.002 |
|  | *Ambispora* | 0.381 | 0.001 |
|  | *Scutellospora* | 0.338 | 0.028 |
| Field and fallow communities | *Glomus* | 0.36 | 0.035 |

VT richness differed significantly between the three root-colonising AM fungal communities (F_2,38_ = 5.91, *P* = 0.005) where the field community had the highest mean richness (8.6 VT ± 0.99) while the fallow community had the lowest (4.2 VT ± 0.46; **Fig. 3a).** Root herbivory had no effect on VT richness (F_2,38_ = 0.31, *P* = 0.58). There were significant dissimilarities in the composition and structure of the three AM fungal communities (PERMANOVA R^2^ = 0.25, F = 6.79, *P* < 0.001; **Fig. 3b),** while root-herbivory had no effect (PERMANOVA R^2^ = 0.015, F = 0.80, *P* = 0.55), and no interaction between factors (PERMANOVA R^2^ = 0.03, F = 0.84, *P* = 0.57). Post-hoc analysis of the AMF factor confirmed all three AM fungal communities were significantly dissimilar from each other **(Fig. 3b; Table S4),** where the forest and fallow AM fungal communities were most dissimilar overall (R^2^=0.28, F = 10.68, *P* < 0.001; **Fig 3b).** Total root colonisation by AM fungi did not differ between the three AM fungal communities, and no colonisation was identified in the no AM fungi plants **(Fig. S1a).**

**Fig. 3.**
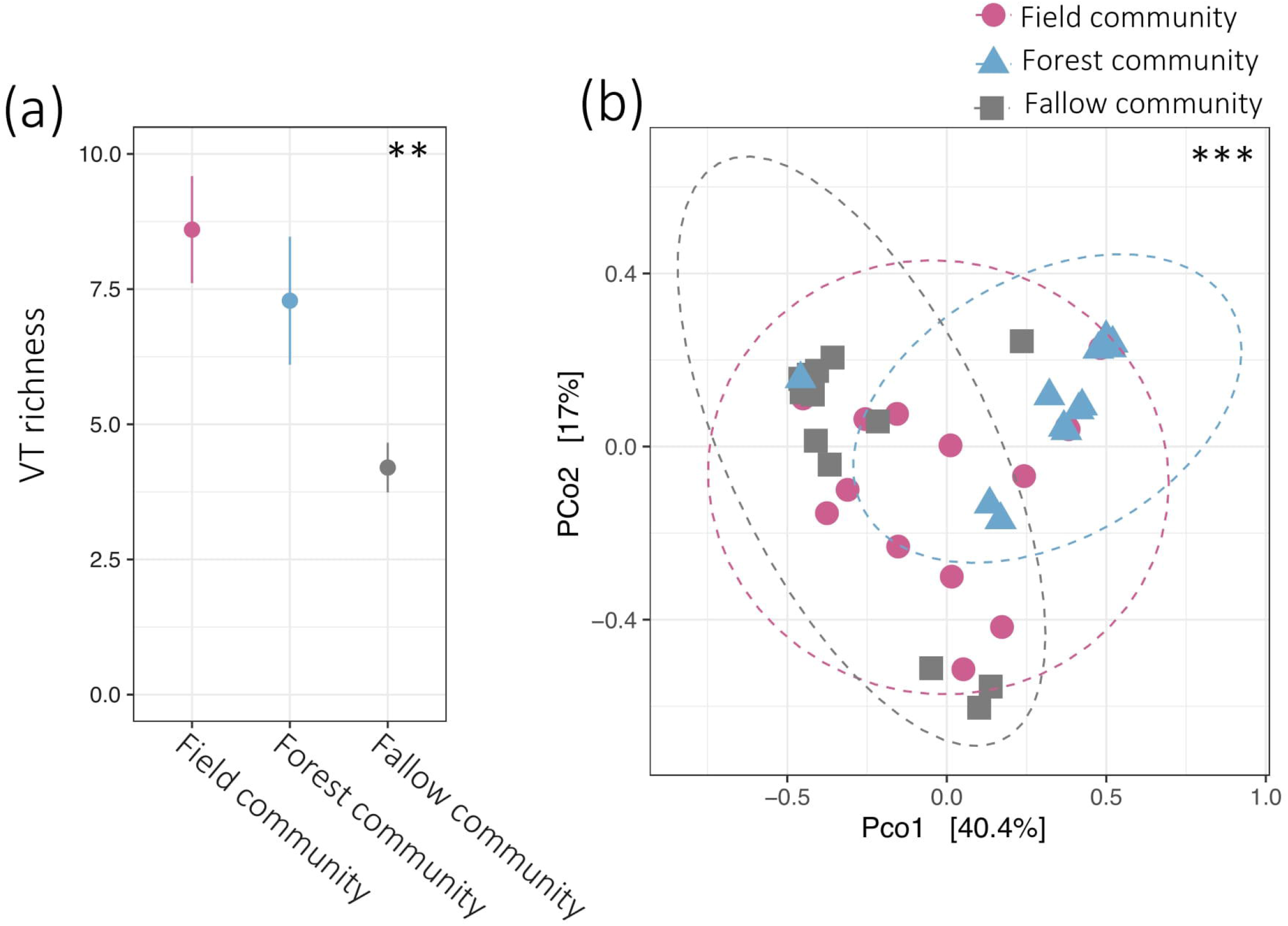
**(a)** Alpha diversity (species richness) of arbuscular mycorrhizal (AM) fungal virtual taxa (VT) in roots colonised by AM fungi sourced from a cropped agricultural field (‘field community’), a sclerophyll forest (‘forest community’), or a cropped agricultural field in fallow (‘fallow community’). Values are means ± SE. **(b)** Principal coordinate analysis of beta diversity (Bray-curtis dissimilarity) comparing the composition and structure of root-colonising AM fungi in the field, forest, and fallow communities. The proportion of the total variation in AM fungal communities explained by the first two principal axes is shown in parentheses. Significant effects of the AM fungal treatments are indicated ** *P* < 0.01.

### Plant biomass and nutrient content

Plant biomass significantly differed between the AM fungal treatments where plants associated with the fallow fungal community had the lowest total biomass, which was 43% lower than the biomass of plants colonised by the forest community **(Fig. 4a; Table S5).** A similar pattern of responses was observed in aboveground and belowground biomass where plants associated with the fallow community had the lowest biomass both above and belowground **(Fig. 4b, c).** Root herbivory significantly reduced total biomass **(Fig. 4a),** an outcome mostly driven by considerable reductions in root biomass in response to herbivory. This effect belowground, however, was dependent on the community of AM fungi, where root-herbivory only reduced root mass in plants with either no AM fungi, or in plants associated with forest community **(Fig. 4c; Table S5).** This AM fungal community-dependent response to root-herbivory was also observed across most root nutrient content responses. Specifically, total root content of phosphorus **(Fig. 5a),** zinc **(Fig. 5b),** manganese **(Fig. 5c),** nitrogen **(Fig. 5d),** magnesium **(Fig. 5e),** sulphur **(Fig. 5f),** and boron **(Fig. 5h)** were reduced by root-herbivory, but only in plants with no AM fungi, or those colonised by the forest fungal community. Although in many cases AM fungi did not increase root nutrient contents in the absence of herbivory, plants associated with the forest community consistently had comparable, or slightly greater, nutrient contents compared to plants without AM fungi. Meanwhile, without any herbivore, plants colonised by the field or fallow communities tended to have lower nutrient contents compared to plants without AM fungi.

**Fig. 4.**
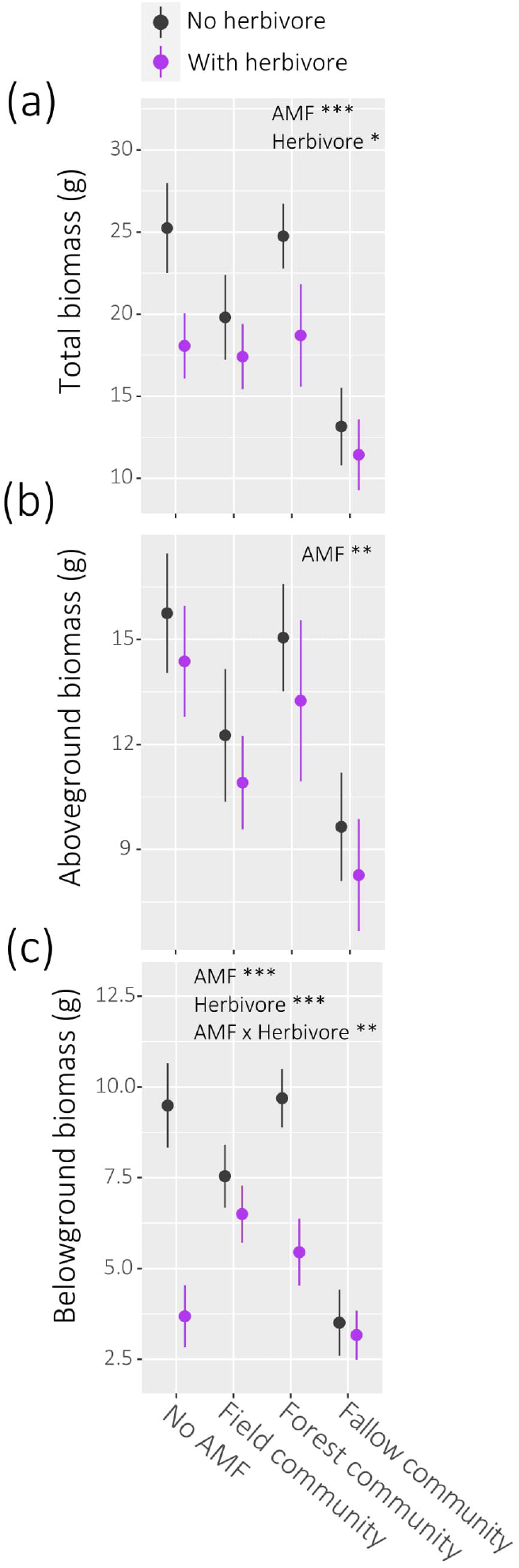
Effects of root-herbivory on the **(a)** total biomass (g), **(b)** aboveground biomass (g), and **(c)** belowground biomass (g) of plants with no arbuscular mycorrhizal (AM) fungi (AMF) and plants associating with one of three different AM fungal communities sourced from a cropped agricultural field (‘field community’), a sclerophyll forest (‘forest community’), or a cropped agricultural field in fallow (‘fallow community’). Significant effects are indicated * *P* <0.05, ** P<0.01, *** *P* <0.001, values are means ± SE.

**Fig. 5.**
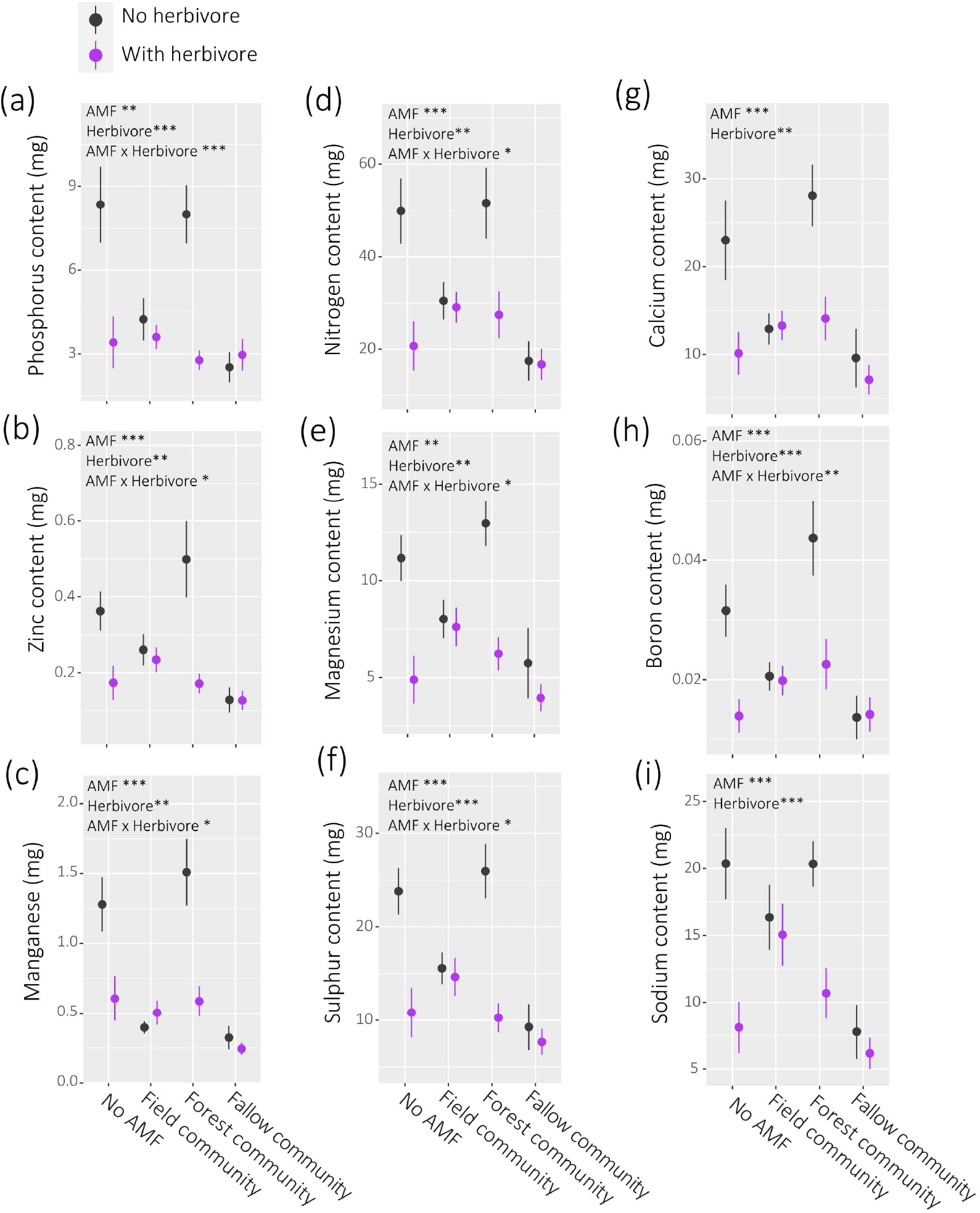
Effects of root-herbivory on the total root content of **(a)** phosphorus (mg), **(b)** zinc (mg), **(c)** manganese (mg), **(d)** nitrogen (mg), **(e)** Magnesium (mg), (f) sulphur, **(g)** calcium (mg), **(h)** boron (mg), and **(i)** sodium in plants with no arbuscular mycorrhizal (AM) fungi (AMF) and in plants associating with one of three different AM fungal communities sourced from a cropped agricultural field (‘field community’), a sclerophyll forest (‘forest community’), or a cropped agricultural field in fallow (‘fallow community’). Significant effects are indicated * *P* <0.05, ** P<0.01, *** *P*

## DISCUSSION

The importance of AM fungi to plant-herbivore interactions is of long-standing interest (Gange, 2001; Gehring & Bennett, 2009; Hartley & Gange, 2009; Koricheva *et al.*, 2009), but progress remains stymied by a narrow focus on a small selection of AM fungal taxa (Malik, 2018; Albornoz *et al.*, 2021; Frew *et al.*, 2022). We investigated how the growth and belowground nutrient status of an important crop species responded to root-herbivory when associating with different field-sourced AM fungal communities.

Plant growth and nutrient responses to root-herbivory were dependent on the community of root-colonising AM fungi. Specifically, when attacked by a root herbivore, plants associating with AM fungal communities sourced from the crop field soil or fallow field soil did not exhibit reductions in root mass, or reductions in root nutrient content. In contrast, when plants with no AM fungi or plants with the forest-derived fungal community were subjected to root-herbivory, they suffered significant losses in root mass and in root content of phosphorus, zinc, manganese, nitrogen, magnesium, sulphur, and boron. Notably, the field fungal community had the highest VT richness while fallow had the lowest, which does not support our first hypothesis that higher alpha diversity would be associated with greater tolerance to herbivory. This highlights that the role of AM fungi in plant responses to root-herbivory cannot be gauged simply by alpha diversity metrics alone. That said, the composition and structure of the field and fallow communities were more similar to each other than to the forest community. This is perhaps unsurprising as the field and fallow AM fungal communities were sourced from fields with histories of horticultural cropping, while the forest community was sourced from soils taken from an adjacent forest habitat.

Both the field and fallow communities had high relative abundances of *Glomus* and *Claroideoglomus.* Furthermore, their indicator fungal VTs were all *Claroideoglomus* spp., while the indicator genera analysis also revealed *Claroideoglomus*, along with *Glomus*, were strongly associated with the fallow community. In contrast to this, the forest community was dominated by the genera *Ambispora* and *Paraglomus*, and had four indicator VTs, all different genera **(Table 1**). This supports our second hypothesis that plant responses to root-herbivory would vary with differences in fungal community composition and structure. It suggests that *Claroideoglomus* spp. may have been more effective at supporting plant tolerance to root-herbivory, where plants dominated by these fungal taxa did not suffer the reductions in growth or nutrients that other plants did when attacked by the root-herbivore. Both *Glomus* and *Claroideoglomus* taxa have previously been associated with a ruderal life history strategy (Hart & Reader, 2002; Chagnon *et al.*, 2013), often found to dominate regularly-disturbed agricultural landscapes (Jansa *et al.*, 2002; Chagnon *et al.*, 2013). As ruderal genera, they may be characterised as fast growing and able to quickly re-establish hyphal networks (Chagnon *et al.*, 2013). Furthermore, they have superior precision and speed when it comes to foraging for nutrients compared with other taxa, including *Paraglomus* spp. (Šmilauer *et al.*, 2020). Therefore, in this instance, such ruderal traits may have allowed *Claroideoglomus* to support plant re-growth and nutrient uptake under root-herbivory, while the fungi comprising the forest community (more dominated by other taxa including *Paraglomus* spp.) were less effective at supporting these tolerance-associated responses to herbivory. It has also been suggested that ruderal AM fungi may be particularly supportive of plant defence responses to pathogens and herbivory (Newsham *et al.*, 1995; Maherali & Klironomos, 2007; Frew *et al.*, 2022) which would potentially correspond with our findings here.

AM fungi that do not have a ruderal life history strategy may be more sensitive to hyphal damage, take longer to recover, and take longer to forage and deliver resources to their host. However, it is important to highlight that there remains little consensus on which AM fungal taxa can be classified as ruderal. Further, as most of our knowledge of AM fungal functional ecology is based on a few well-studied taxon groups, particularly Glomerales, it is difficult to draw general conclusions about the roles in different contexts. Compared with *Glomus* and *Claroideoglomus*, notably less is known about the life history strategies of *Ambispora* and *Paraglomus*, which were more abundant in the forest-derived fungal community. Taxa from these genera are indeed often detected in non-agricultural systems, including forests and unmanaged woodlands (Chagnon *et al.*, 2013; Prober *et al.*, 2015; De Beenhouwer *et al.*, 2015). *Paraglomus* has been shown to be sensitive to different types of disturbance, particularly high intensity agriculture (Hijri *et al.*, 2006; Gosling *et al.*, 2014). Therefore, if these AM fungi were not as quick or effective at establishing their hyphal networks and foraging for nutrients, this may explain why plants colonised by the forest community were more susceptible to the negative outcomes of root-herbivory.

That said, different plant outcomes might have been observed if more time was given in this experiment where other fungal taxa (e.g., *Paraglomus* and *Ambispora*) may have more opportunity to establish, to forage for nutrients and perhaps provide greater support for plants under herbivory. Yet, both *Paraglomus* and *Ambispora* have also been found to be positively associated with habitats with high grazing (Heyde *et al.*, 2017), which would not support the theory they are highly-sensitive to disturbance. Fungal relatedness in our three communities may also have affected the amount of ‘benefit’ plants could derive from their AM fungi when dealing with herbivore attack. Specifically, the field and fallow communities were significantly dominated by Glomerales (>75%), while the forest fungal communities had greater abundances of other orders (Paraglomerales, Archaeosporales, Diversisporales). Studies suggest closely related fungal communities can offer better support for plant growth and nutrient acquisition, compared to less related fungal networks (Roger *et al.*, 2013; van’t Padje *et al.*, 2022), perhaps due to reduced competition. However, plants can also be expected to benefit from associating with unrelated fungi where there is variation in their abilities to acquire different resources or provide other functions (Argüello *et al.*, 2016). It is clear we need to broaden our understanding of the ecology and functional traits of different AM fungal taxa (Van Der Heijden & Scheublin, 2007; Chaudhary *et al.*, 2016) if we are to elucidate their functional significance in plant-herbivore interactions.

It is notable that the three AM fungal communities provided little or no ‘benefit’ to their hosts in terms of biomass or nutrient uptake in the absence of herbivory. In these instances, the outcomes of associating with AM fungi may be interpreted as negative overall, compared with plants with no AM fungi. For example, the total biomass of plants without root herbivory was lower when associating with field or fallow fungal communities, compared with plants with no AM fungi. A similar pattern was observed for root nutrient status, where the field and fallow communities appear to have reduced root nutrient content in plants without herbivory. However, these two fungal communities did provide benefits to their hosts under herbivory pressure. Contrastingly, plants with the forest-sourced fungi displayed significant biomass and nutrient losses under herbivory, yet these fungi did provide nutrient benefits (e.g., zinc, manganese, boron) compared to plants without AM fungi. As mentioned previously, this may be associated with the amount of cooperation and competition between fungal taxa within our communities (van’t Padje *et al.*, 2022), and it is possible the results would have been different if more time was given for the fungal communities to develop further. Particular functional ‘benefits’ that AM fungi provide to their hosts may not necessarily be apparent unless faced with particular environmental conditions (Veresoglou *et al.*, 2021).

While AM fungi are commonly characterised by their ability to enhance plant nutrient acquisition and growth, there are many reported instances where they do not generate these particular outcomes for their hosts (Johnson *et al.*, 1997; Klironomos, 2003; Jacott *et al.*, 2017). It is increasingly acknowledged that this is a very narrow understanding of AM fungal ecology, a dogma that is perhaps driven by long-standing research bias towards particular functions (e.g. increased plant growth, phosphorus uptake) and particular plant and fungal taxa (Chaudhary *et al.*, 2016; Albornoz *et al.*, 2021), including in the context of plant-herbivore interactions (Malik, 2018; Frew *et al.*, 2022). Albornoz *et al.* (2021) recently emphasised the need for a greater appreciation of the diverse functions and benefits of AM fungi (Delavaux *et al.*, 2017) and to better understand drivers of their context dependency.

Our findings indicate that the growth and belowground nutrient responses of a major crop species to root-herbivory differs depending on the composition and structure of the root-colonising AM fungal community. In this context, plants associating with fungal communities dominated by *Glomus* and *Claroideoglomus* were less susceptible to herbivory-induced losses in belowground biomass and nutrients, while colonisation by communities with high abundances of *Paraglomus* and *Ambispora* were associated with growth and nutrient reductions. This study did not seek to ascertain the roles and effects of specific fungal taxa in isolation but it is important for the research community to increase exploration of how natural assemblages of AM fungi are associated with plant responses to herbivory. This is an important step in advancing our ability to effectively manage local AM fungal communities to support soil sustainability in agriculture.

## ACKNOWLEDGEMENTS

The authors would like to thank the Institute for Life Sciences and the Environment, the Centre for Crop Health, and the Next Generation Sequencing Facility at Western Sydney University for their technical assistance and support. This work was supported by a Thomas Davies Research Grant for Marine, Soil and Plant Biology administered by the Australian Academy of Science, awarded to AF. AF was supported by an Australian Research Council Discovery Early Career Researcher Award (DE220100479). The authors would like to thank Dr Pauline Lenancker from Sugar Research Australia Pty Ltd for supplying *D. albohirtum* larvae.

## AUTHORS’ CONTRIBUTIONS

All authors contributed to the study conception and design. AF and AN established the experiment; AN led sample processing along with AF and BALW; AF and AN conducted statistical analysis and all authors contributed to the interpretation of the results. The first manuscript draft was led by AN as part of a Master of Science thesis, all authors then contributed to final manuscript development.

## DATA AVAILABILITY

Data that support this study are available in the figshare repository at the following DOI: 10.6084/m9.figshare.19166444. Raw DNA sequencing data are available under NCBI BioProject accession number PRJNA805065.

## Notes

### Competing Interest Statement

The authors have declared no competing interest.

## REFERENCES

Albornoz FE, Dixon KW, Lambers H. 2021. Revisiting mycorrhizal dogmas: Are mycorrhizas really functioning as they are widely believed to do? Soil Ecology Letters 3: 73–82.

Allsopp PG. 2010. Integrated management of sugarcane whitegrubs in Australia: an evolving success. Annual Review of Entomology 55: 329–349.

Argüello A, O’Brien MJ, van der Heijden MGA, Wiemken A, Schmid B, Niklaus PA. 2016. Options of partners improve carbon for phosphorus trade in the arbuscular mycorrhizal mutualism. Ecology Letters 19: 648–656.

Bainard LD, Bainard JD, Hamel C, Gan Y. 2014. Spatial and temporal structuring of arbuscular mycorrhizal communities is differentially influenced by abiotic factors and host crop in a semi-arid prairie agroecosystem. FEMS Microbiology Ecology 88: 333–344.

Barber NA, Kiers ET, Theis N, Hazzard RV, Adler LS. 2013. Linking agricultural practices, mycorrhizal fungi, and traits mediating plant–insect interactions. Ecological Applications 23: 1519–1530.

Bardgett RD, van der Putten WH. 2014. Belowground biodiversity and ecosystem functioning. Nature 515: 505–511.

Bender SF, Wagg C, van der Heijden MGA. 2016. An underground revolution: Biodiversity and soil ecological engineering for agricultural sustainability. Trends in Ecology & Evolution 31: 440–452.

Bennett AE, Bever JD. 2007. Mycorrhizal species differentially alter plant growth and response to herbivory. Ecology 88: 210–218.

Bennett AE, Bever JD, Bowers MD. 2009. Arbuscular mycorrhizal fungal species suppress inducible plant responses and alter defensive strategies following herbivory. Oecologia 160: 771–779.

Borowicz VA. 2013. The impact of arbuscular mycorrhizal fungi on plant growth following herbivory: A search for pattern. Acta Oecologica 52: 1–9.

Cáceres MD, Legendre P. 2009. Associations between species and groups of sites: indices and statistical inference. Ecology 90: 3566–3574.

Caporaso JG, Ackermann G, Apprill A, Bauer M, Berg-Lyons D, Betley J, Fierer N, Fraser L, Fuhrman JA, Gilbert JA, et al. 2018. EMP 16S Illumina amplicon protocol. Protocols.io: See http://www.earthmicrobiome.org/protocols-and-standards/16s.

Chagnon P-L, Bradley RL, Maherali H, Klironomos JN. 2013. A trait-based framework to understand life history of mycorrhizal fungi. Trends in Plant Science 18: 484–491.

Chaudhary VB, Rúa MA, Antoninka A, Bever JD, Cannon J, Craig A, Duchicela J, Frame A, Gardes M, Gehring C, et al. 2016. MycoDB, a global database of plant response to mycorrhizal fungi. Scientific Data 3: 160028.

De Beenhouwer M, Van Geel M, Ceulemans T, Muleta D, Lievens B, Honnay O. 2015. Changing soil characteristics alter the arbuscular mycorrhizal fungi communities of Arabica coffee (Coffea arabica) in Ethiopia across a management intensity gradient. Soil Biology and Biochemistry 91: 133–139.

Delavaux CS, Smith-Ramesh LM, Kuebbing SE. 2017. Beyond nutrients: a meta-analysis of the diverse effects of arbuscular mycorrhizal fungi on plants and soils. Ecology 98: 2111–2119.

Dumbrell AJ, Ashton PD, Aziz N, Feng G, Nelson M, Dytham C, Fitter AH, Helgason T. 2011. Distinct seasonal assemblages of arbuscular mycorrhizal fungi revealed by massively parallel pyrosequencing. New Phytologist 190: 794–804.

Fox J, Weisberg S. 2011. An R Companion to Applied Regression. Thousand Oaks, CA: Sage Publications.

Frew A, Antunes PM, Cameron DD, Hartley SE, Johnson SN, Rillig MC, Bennett AE. 2022. Plant herbivore protection by arbuscular mycorrhizas: a role for fungal diversity? New Phytologist 233: 1022–1031.

Frew A, Powell JR, Allsopp PG, Sallam N, Johnson SN. 2017. Arbuscular mycorrhizal fungi promote silicon accumulation in plant roots, reducing the impacts of root herbivory. Plant and Soil 419: 423–433.

Frew A, Powell JR, Johnson SN. 2020. Aboveground resource allocation in response to root herbivory as affected by the arbuscular mycorrhizal symbiosis. Plant and Soil 447: 463–473.

Frew A, Wilson BAL. 2021. Different mycorrhizal fungal communities differentially affect plant phenolic-based resistance to insect herbivory. Rhizosphere 19: 100365.

Gange AC. 2001. Species-specific responses of a root-and shoot-feeding insect to arbuscular mycorrhizal colonization of its host plant. New Phytologist 150: 611–618.

Gehring C, Bennett A. 2009. Mycorrhizal fungal-plant-insect interactions: the importance of a community approach. Environmental Entomology 38: 93–102.

Gosling P, Proctor M, Jones J, Bending GD. 2014. Distribution and diversity of Paraglomus spp. in tilled agricultural soils. Mycorrhiza 24: 1–11.

Hart MM, Antunes PM, Chaudhary VB, Abbott LK. 2018. Fungal inoculants in the field: Is the reward greater than the risk? Functional Ecology 32: 126–135.

Hart MM, Reader RJ. 2002. Taxonomic basis for variation in the colonization strategy of arbuscular mycorrhizal fungi. New Phytologist 153: 335–344.

Hartley SE, Gange AC. 2009. Impacts of plant symbiotic fungi on insect herbivores: mutualism in a multitrophic context. Annual Review of Entomology 54: 323–342.

He L, Li C, Liu R. 2017. Indirect interactions between arbuscular mycorrhizal fungi and *Spodoptera exigua* alter photosynthesis and plant endogenous hormones. Mycorrhiza 27: 525–535.

Hervé M, Hervé MM. 2020. Package ‘RVAideMemoire’. See https://CRAN.R-project.org/package=RVAideMemoire.

Heyde M van der, Bennett JA, Pither J, Hart M. 2017. Longterm effects of grazing on arbuscular mycorrhizal fungi. Agriculture, Ecosystems & Environment 243: 27–33.

Hijri I, Sýkorová Z, Oehl F, Ineichen K, Mäder P, Wiemken A, Redecker D. 2006. Communities of arbuscular mycorrhizal fungi in arable soils are not necessarily low in diversity. Molecular Ecology 15: 2277–2289.

Hill EM, Robinson LA, Abdul-Sada A, Vanbergen AJ, Hodge A, Hartley SE. 2018. Arbuscular mycorrhizal fungi and plant chemical defence: effects of colonisation on aboveground and belowground metabolomes. Journal of Chemical Ecology 44: 198–208.

Jacott CN, Murray JD, Ridout CJ. 2017. Trade-offs in arbuscular mycorrhizal symbiosis: Disease resistance, growth responses and perspectives for crop breeding. Agronomy 7: 75.

Jansa J, Mozafar A, Anken T, Ruh R, Sanders I, Frossard E. 2002. Diversity and structure of AMF communities as affected by tillage in a temperate soil. Mycorrhiza 12: 225–234.

Johnson NC, Graham J-H, Smith FA. 1997. Functioning of mycorrhizal associations along the mutualism–parasitism continuum. New Phytologist 135: 575–585.

Johnson SN, Rasmann S. 2015. Root-feeding insects and their interactions with organisms in the rhizosphere. Annual Review of Entomology 60: 517–535.

Klironomos JN. 2003. Variation in plant response to native and exotic arbuscular mycorrhizal fungi. Ecology 84: 2292–2301.

Koide RT, Li M. 1989. Appropriate controls for vesicular–arbuscular mycorrhiza research. New Phytologist 111: 35–44.

Koricheva J, Gange AC, Jones T. 2009. Effects of mycorrhizal fungi on insect herbivores: a meta-analysis. Ecology 90: 2088–2097.

Kula AAR, Hartnett DC, Wilson GWT. 2005. Effects of mycorrhizal symbiosis on tallgrass prairie plant–herbivore interactions. Ecology Letters 8: 61–69.

Lee J, Lee S, Young JPW. 2008. Improved PCR primers for the detection and identification of arbuscular mycorrhizal fungi. FEMS Microbiology Ecology 65: 339–349.

Maherali H, Klironomos JN. 2007. Influence of phylogeny on fungal community assembly and ecosystem functioning. Science 316: 1746–1748.

Malik RJ. 2018. Recent trend: Is the role of arbuscular mycorrhizal fungi in plant-enemies performance biased by taxon usage? The American Midland Naturalist 180: 306–311.

McGonigle TP, Miller MH, Evans DG, Fairchild GL, Swan JA. 1990. A new method which gives an objective measure of colonization of roots by vesicular—arbuscular mycorrhizal fungi. New Phytologist 115: 495–501.

McMurdie PJ, Holmes S. 2013. phyloseq: An R Package for Reproducible Interactive Analysis and Graphics of Microbiome Census Data. PLOS ONE 8: e61217.

Moora M, Davison J, Öpik M, Metsis M, Saks Ü, Jairus T, Vasar M, Zobel M. 2014. Anthropogenic land use shapes the composition and phylogenetic structure of soil arbuscular mycorrhizal fungal communities. FEMS Microbiology Ecology 90: 609–621.

Newsham KK, Fitter AH, Watkinson AR. 1995. Multi-functionality and biodiversity in arbuscular mycorrhizas. Trends in Ecology & Evolution 10: 407–411.

Oksanen J, Blanchet FG, Legendre P, Minchin PR, O’Hara RB, Simpson GL, Solymos P, Stevens MHH, Wagner H. 2015. vegan: Community Ecology Package.: http://CRAN.R-project.org/package=vegan.

Öpik M, Davison J. 2016. Uniting species- and community-oriented approaches to understand arbuscular mycorrhizal fungal diversity. Fungal Ecology 24: 106–113.

Öpik M, Vanatoa A, Vanatoa E, Moora M, Davison J, Kalwij JM, Reier Ü, Zobel M. 2010. The online database MaarjAM reveals global and ecosystemic distribution patterns in arbuscular mycorrhizal fungi (Glomeromycota). New Phytologist 188: 223–241.

Öpik M, Zobel M, Cantero JJ, Davison J, Facelli JM, Hiiesalu I, Jairus T, Kalwij JM, Koorem K, Leal ME, et al. 2013. Global sampling of plant roots expands the described molecular diversity of arbuscular mycorrhizal fungi. Mycorrhiza 23: 411–430.

van’t Padje A, Klein M, Caldas V, Oyarte Galvez L, Broersma C, Hoebe N, Sanders IR, Shimizu T, Kiers ET. 2022. Decreasing relatedness among mycorrhizal fungi in a shared plant network increases fungal network size but not plant benefit. Ecology Letters 25: 509–520.

Powell JR, Rillig MC. 2018. Biodiversity of arbuscular mycorrhizal fungi and ecosystem function. New Phytologist 220: 1059–1075.

Prober SM, Bissett A, Walker C, Wiehl G, McIntyre S, Tibbett M. 2015. Spatial structuring of arbuscular mycorrhizal communities in benchmark and modified temperate eucalypt woodlands. Mycorrhiza 25: 41–54.

R Core Team. 2017. R: A Language and Environment for Statistical Computing. Vienna, Austria: R Foundation for Statistical Computing.

Rayment GE, Lyons DJ. 2011. Soil chemical methods: Australasia. Melbourne, Australia: CSIRO publishing.

Rillig MC, Aguilar-Trigueros CA, Camenzind T, Cavagnaro TR, Degrune F, Hohmann P, Lammel DR, Mansour I, Roy J, van der Heijden MGA, et al. 2019. Why farmers should manage the arbuscular mycorrhizal symbiosis. New Phytologist 222: 1171–1175.

Rodriguez A, Sanders IR. 2015. The role of community and population ecology in applying mycorrhizal fungi for improved food security. The ISME Journal 9: 1053–1061.

Roger A, Colard A, Angelard C, Sanders IR. 2013. Relatedness among arbuscular mycorrhizal fungi drives plant growth and intraspecific fungal coexistence. The ISME Journal 7: 2137–2146.

Rognes T, Flouri T, Nichols B, Quince C, Mahé F. 2016. VSEARCH: a versatile open source tool for metagenomics. PeerJ 4: e2584.

Sallam N. 2011. Review of current knowledge on the population dynamics of *Dermolepida albohirtum* (Waterhouse)(Coleoptera: Scarabaeidae). Australian Journal of Entomology 50: 300–308.

Slansky FJ. 1985. Food utilization by insects: Interpretation of observed differences between dry weight and energy efficiencies. Entomologia Experimentalis et Applicata 39: 47–60.

Šmilauer P, Šmilauerová M, Kotilínek M, Košnar J. 2020. Foraging speed and precision of arbuscular mycorrhizal fungi under field conditions: An experimental approach. Molecular Ecology 29: 1574–1587.

Smith SE, Read DJ. 2008. Mycorrhizal Symbiosis. Amsterdam, the Netherlands & Boston, MA: Academic Press.

Spatafora JW, Chang Y, Benny GL, Lazarus K, Smith ME, Berbee ML, Bonito G, Corradi N, Grigoriev I, Gryganskyi A, et al. 2016. A phylum-level phylogenetic classification of zygomycete fungi based on genome-scale data. Mycologia 108: 1028–1046.

Tao L, Ahmad A, Roode JC de, Hunter MD. 2016. Arbuscular mycorrhizal fungi affect plant tolerance and chemical defences to herbivory through different mechanisms. Journal of Ecology 104: 561–571.

Tedersoo L, Bahram M, Zobel M. 2020. How mycorrhizal associations drive plant population and community biology. Science 367: eaba1223.

Thirkell TJ, Charters MD, Elliott AJ, Sait SM, Field KJ. 2017. Are mycorrhizal fungi our sustainable saviours? Considerations for achieving food security. Journal of Ecology 105: 921–929.

Van Der Heijden MG, Scheublin TR. 2007. Functional traits in mycorrhizal ecology: their use for predicting the impact of arbuscular mycorrhizal fungal communities on plant growth and ecosystem functioning. New Phytologist 174: 244–250.

Vannette RL, Hunter MD, Rasmann S. 2013. Arbuscular mycorrhizal fungi alter above- and below-ground chemical defense expression differentially among *Asclepias* species. Frontiers in Plant Science 4: 361.

Vannette RL, Rasmann S. 2012. Arbuscular mycorrhizal fungi mediate below-ground plant–herbivore interactions: a phylogenetic study. Functional Ecology 26: 1033–1042.

Vasar M, Andreson R, Davison J, Jairus T, Moora M, Remm M, Young JPW, Zobel M, Öpik M. 2017. Increased sequencing depth does not increase captured diversity of arbuscular mycorrhizal fungi. Mycorrhiza 27: 761–773.

Vasar M, Davison J, Neuenkamp L, Sepp S-K, Young JPW, Moora M, Öpik M. 2021. User-friendly bioinformatics pipeline gDAT (graphical downstream analysis tool) for analysing rDN A sequences. Molecular Ecology Resources.

Venables WN, Ripley BD. 2002. Modern Applied Statistics with S. New York: Springer.

Veresoglou SD, Johnson D, Mola M, Yang G, Rillig MC. 2021. Evolutionary bet-hedging in arbuscular mycorrhiza-associating angiosperms. New Phytologist: 10.1111/nph.17852.

Vierheilig H, Coughlan AP, Wyss U, Piché Y. 1998. Ink and vinegar, a simple staining technique for arbuscular-mycorrhizal fungi. Applied and Environmental Microbiology 64: 5004–5007.

Wooley SC, Paine TD. 2007. Can intra-specific genetic variation in arbuscular mycorrhizal fungi (*Glomus etunicatum*) affect a mesophyll-feeding herbivore (*Tupiocoris notatus* Distant)? Ecological Entomology 32: 428–434.

Zuur AF, Ieno EN, Elphick CS. 2010. A protocol for data exploration to avoid common statistical problems. Methods in Ecology and Evolution 1: 3–14.

