## Supplementary material for "Belowground plant responses to root herbivory depend on the composition and structure of their root-colonising arbuscular mycorrhizal fungi": Table S1; Table S2; Table S3; Table S4; Table S5; Fig. S1

**TABLE S1.** Soil substrate details. Analysis of homogenised growth substrate in pots under different AMF soil treatments. Analysis carried out by CSBP Soil & Plant Analysis Laboratory, Bibra Lake, WA, Australia.

|  | Nitrogen (ammonium) (mg/kg) | Phosphorus (colwell) (mg/kg) | Potassium (colwell) (mg/kg) | Sulphur (mg/kg) | Organic carbon (%) | pH |
| --- | --- | --- | --- | --- | --- | --- |
| Field soil | 2.01 ± 0.19 | 72.33 ± 9.25 | 356 ± 31.88 | 148.17 ± 4.05 | 2.75 ± 0.55 | 5.73 ± 0.18 |
| Forest soil | 2.33 ± 0.33 | 69.33 ± 6.56 | 335 ± 21.67 | 145 ± 2.98 | 2.25 ± 0.18 | 5.67 ± 0.03 |
| Fallow soil | 2.11 ± 0.01 | 68.67 ± 7.69 | 358 ± 42.19 | 143.67 ± 2.39 | 2.29 ± 0.54 | 5.83 ± 0.03 |
| No AMF | 2.67 ± 0.34 | 67.88 ± 5.24 | 339 ± 22.12 | 145.9 ± 4.24 | 2.32 ± 0.14 | 5.87 ± 0.33 |

**Table S2**. List of virtual taxa (VT) of arbuscular mycorrhizal (AM) fungi showing taxonomy referenced with Maarj*AM* (≥ 97% match).

| AM fungal VT ID | Taxonomy |
| --- | --- |
| VTX00193 | Claroideoglomeraceae, Claroideoglomus lamellosum T VTX00193 |
| VTX00444 | Paraglomeraceae, Paraglomus IH1 VTX00444 |
| VTX00067 | Glomeraceae, Glomus mosseae VTX00067 |
| VTX00092 | Glomeraceae ,Glomus sp. VTX00092 |
| VTX00242 | Ambisporaceae, Ambispora leptoticha T VTX00242 |
| VTX00057 | Claroideoglomeraceae, Claroideoglomus sp. VTX00057 |
| VTX00238 | Paraglomeraceae, Paraglomus occultum T VTX00238 |
| VTX00111 | Glomeraceae, Glomus Afrothismia foertheriana symbiont T VTX00111 |
| VTX00245 | Archaeosporaceae, Archaeospora Shi14b Arc-27 VTX00245 |
| VTX00005 | Archaeosporaceae, Archaeospora sp. VTX00005 |
| VTX00228 | Acaulosporaceae, Acaulospora sp. VTX00228 |
| VTX00191 | Glomeraceae, Glomus A VTX00191 |
| VTX00052 | Gigasporaceae, Scutellospora aurigloba VTX00052 |
| VTX00213 | Glomeraceae, Glomus sp. T VTX00213 |
| VTX00074 | Glomeraceae, Glomus sp. VTX00074 |
| VTX00030 | Acaulosporaceae, Acaulospora MO-A9 VTX00030 |
| VTX00327 | Glomeraceae, Glomus sp. T VTX00327 |
| VTX00065 | Glomeraceae, Glomus sp. VTX00065 |
| VTX00338 | Archaeosporaceae, Archaeospora sp. VTX00338 |
| VTX00279 | Claroideoglomeraceae, Claroideoglomus sp. T VTX00279 |
| VTX00340 | Claroideoglomeraceae, Claroideoglomus Glo G8 VTX00340 |
| VTX00049 | Gigasporaceae, Scutellospora sp. VTX00049 |
| VTX00056 | Claroideoglomeraceae, Claroideoglomus Alguacil12b GLO G1 VTX00056 |
| VTX00060 | Diversisporaceae, Diversispora celata VTX00060 |
| VTX00009 | Archaeosporaceae, Archaeospora sp. VTX00009 |
| VTX00023 | Acaulosporaceae, Acaulospora sp. VTX00023 |
| VTX00219 | Glomeraceae, Glomus sp. VTX00219 |
| VTX00239 | Paraglomeraceae, Paraglomus sp. VTX00239 |
| VTX00375 | Paraglomeraceae, Paraglomus MO-P1 VTX00375 |
| VTX00024 | Acaulosporaceae, Acaulospora lacunosa VTX00024 |
| VTX00039 | Gigasporaceae, Gigaspora margarita VTX00039 |
| VTX00113 | Glomeraceae, Glomus sp. VTX00113 |
| VTX00433 | Paraglomeraceae, Paraglomus MO-P2 VTX00433 |
| VTX00446 | Paraglomeraceae, Paraglomus IS-Pg1 VTX00446 |

**Table S3**. Results of generalised linear models examining effects of ‘AMF’ factor on relative sequence abundance of AM fungal genera.

|  | F (Chisq) | *P* |
| --- | --- | --- |
| *Glomus* | 8.3196 | 0.016 |
| *Claroideoglomus* | 13.97 | <0.001 |
| *Paraglomus* | 12.292 | 0.002 |
| *Ambispora* | 20.907 | <0.001 |
| *Acaulospora* | 10.66 | 0.005 |
| *Archaeospora* | 5.073 | 0.079 |
| *Scutellospora* | 18.086 | <0.001 |
| *Diversispora* | 6.281 | 0.05 |
| *Gigaspora* | 7.095 | 0.029 |

**Table S4**. Results of pairwise post-hoc analysis of PERMANOVA examining differences (Bray Curtis dissimilarity) in composition and structure of the three root-colonising AM fungal communities.

|  | F | *P* | R^2^ |
| --- | --- | --- | --- |
| Field community – Fallow community | 4.14 | 0.006 | 0.128 |
| Forest community – Fallow community | 10.68 | <0.001 | 0.284 |
| Field community – Forest community | 5.61 | <0.001 | 0.172 |

**Table S5**. Results of ANOVAs on fitted standard linear models examining main effects and interaction of factors ‘AMF’ and ‘Herbivore’ on plant biomass and root nutrient content. Results also shown of generalised linear model assessing effects on root colonisation by arbuscular mycorrhizal (AM) fungi, and results of ANOVAs examining effects of the AMF factors on relative growth rate of the root-herbivore (*Dermolepida albohirtum*).

|  | AMF | | Herbivore | | AMF x Herbivore | |
| --- | --- | --- | --- | --- | --- | --- |
|  | F _3,56_ | *P* | F _3,56_ | *P* | F _3,56_ | *P* |
| Total biomass | **6.812** | **<0.001** | **6.528** | **0.013** | 0.625 | 0.602 |
| Aboveground biomass | **5.173** | **0.003** | 1.492 | 0.227 | 0.008 | 0.999 |
| Belowground biomass | **9.301** | **<0.001** | **20.959** | **<0.001** | **4.321** | **0.008** |
| Phosphorus * | **6.197** | **0.001** | **16.037** | **<0.001** | **6.407** | **<0.001** |
| Zinc * | **8.521** | **<0.001** | **10.796** | **0.002** | **3.718** | **0.017** |
| Manganese * | **17.462** | **<0.001** | **10.607** | **0.002** | **3.986** | **0.012** |
| Nitrogen * | **8.58** | **<0.001** | **10.145** | **0.002** | **3.293** | **0.027** |
| Magnesium | **5.836** | **0.002** | **21.472** | **<0.001** | **3.739** | **0.016** |
| Sulphur * | **8.255** | **<0.001** | **16.057** | **<0.001** | **3.806** | **0.015** |
| Calcium * | **8.801** | **<0.001** | **8.74** | **0.005** | 2.178 | 0.101 |
| Boron | **8.725** | **<0.001** | **12.969** | **<0.001** | **4.295** | **0.008** |
| Sodium | **9.411** | **<0.001** | **13.021** | **<0.001** | 2.579 | 0.063 |
|  | F _3,56_ | *P* (Chisq) | F _3,56_ | *P* (Chisq) | F _3,56_ | *P* (Chisq) |
| Total colonisation † | **88.071** | **<0.001** | 0.016 | 0.898 | 3.842 | 0.279 |
|  | F _3,24_ | *P* |  |  |  |  |
| Herbivore relative growth rate | 0.152 | 0.928 |  |  |  |  |

* Log transformed

† Generalised linear model

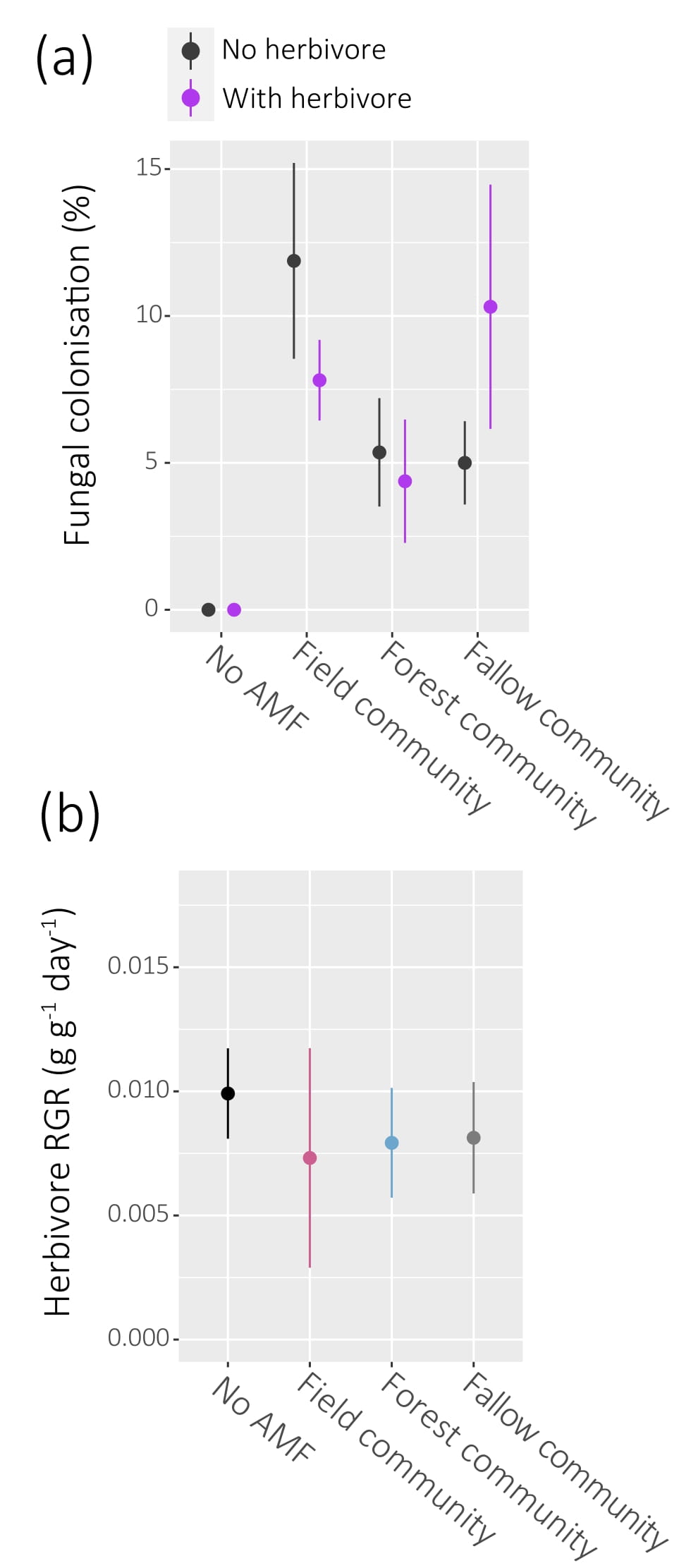

**Figure S1 (a)** Effects of herbivory on total fungal colonisation (%) by arbuscular mycorrhizal (AM) fungi on roots of *Sorghum bicolor* colonised by different field-sourced AM fungal communities, or with no AM fungi (no AMF). **(b)** The relative growth rates of root-herbivores (*Dermolepida albohirtum*) feeding on *Sorghum bicolor* colonised by different field-sources AM fungal communities, or with no AM fungi.
